# The effects of dyslipidaemia and cholesterol modulation on erythrocyte susceptibility to malaria parasite infection

**DOI:** 10.1101/630251

**Authors:** Marion Koch, Jaimini Cegla, Ben Jones, Yuning Lu, Ziad Mallat, Andrew Blagborough, Fiona Angrisano, Jake Baum

**Author notes:** **Corresponding Authors:** Jake Baum. Department of Life Sciences, Imperial College London, Exhibition Road, South Kensington, London, SW7 2AZ, UK.

## Abstract

Malaria disease commences when blood-stage parasites, called merozoites, invade human red blood cells (RBCs). Whilst the process of invasion is traditionally seen as being entirely merozoite-driven, emerging data suggests RBC biophysical properties markedly influence invasion. Cholesterol is a major determinant of cell membrane biophysical properties. We set out to assess whether cholesterol content in the RBC membrane affects susceptibility to merozoite invasion. Here we demonstrate that RBC bending modulus (a measure of deformability) is markedly affected by artificial modulation of cholesterol content and negatively correlated with merozoite invasion efficiency. Contextualising this observation, we tested a mouse model of hypercholesterolemia and human clinical samples from patients with a range of serum cholesterol concentrations for parasite susceptibility. Hypercholesterolaemia in both human and murine subjects had little effect merozoite invasion efficiency. Furthermore, on testing, RBC cholesterol content in both murine and human hypercholesterolaemia settings was found to be unchanged from normal controls. Serum cholesterol is, therefore, unlikely to impact on RBC susceptibility to merozoite entry. Our work, however, suggests that native polymorphisms that affect RBC membrane lipid composition would be expected to affect parasite entry. This supports investigation of RBC biophysical properties in endemic settings, which may yet identify naturally protective lipid-related polymorphisms.

## INTRODUCTION

Cholesterol is a key constituent of human cells and plays a key role in modulating membrane properties, influencing both membrane fluidity (Chabanel, *et al* 1983) and stiffness (Henriksen, *et al* 2004, Meleard, *et al* 1997). The effect on these cellular properties is mediated by cholesterol’s flat rigid structure which is defined by the planar tetracyclic ring shape of the molecule. While an elevation in the cholesterol content contained within cell membranes is expected to lead to a reduction in cell elasticity, how these levels are regulated and how dynamic they are in human red blood cells remains largely unknown. A significant change in plasma lipid levels is medically described as dyslipidaemia, a term used to categorize a number of conditions, including hypercholesterolaemia, which is associated with an increase in plasma cholesterol levels. Importantly, whether and how such an increase in plasma cholesterol would affect cellular membrane composition is not clear. For example, a number of studies have found differences in how treatment of elevated plasma cholesterol or clinical conditions with dyslipidaemia is associated with differences in erythrocyte membrane cholesterol levels (Cazzola, *et al* 2011, Martinez, *et al* 1996, Tziakas, *et al* 2007).

Erythrocyte infection lies at the heart of all symptoms of malaria disease. It is established when the blood-stage merozoite form of the *Plasmodium* parasite attaches to and penetrates the erythrocyte with concomitant formation of a parasitophorous vacuole inside (White, *et al* 2014). There is a detailed appreciation of the stepwise molecular events that characterise merozoite invasion (Cowman, *et al* 2012), however, the role the erythrocyte plays in the process was, up until recently, largely overlooked (Koch and Baum 2016). There has been, however, a growing appreciation in the past few years that parasite binding to the erythrocyte, stimulates biophysical changes in the red cell that likely facilitate entry making it energetically more favourable (Koch, *et al* 2017, Sisquella, *et al* 2017). Further, several key polymorphisms that protect against malaria infection may do so directly by modulating erythrocyte biophysical properties (Genton, *et al* 1995, Ndila, *et al* 2018). These polymorphisms are generally associated with changes in either the erythrocyte cytoskeleton (Genton, *et al* 1995) or membrane surface proteins, such as components of the glycophorin family, a well-studied group of erythrocyte surface receptors that are known to be under natural selection, likely from malaria (Baum, *et al* 2002, Malaria Genomic Epidemiology, *et al* 2015, Ndila, *et al* 2018). To date, however, there is little study of the effects of lipid changes in mediating susceptibility to invasion and whether changes in erythrocyte membrane lipid composition might be associated with changes in efficiency of malaria parasite entry. Several studies have explored the complex relationship between obesity, nutrition and malaria. Obesity has been implicated in being protective against cerebral malaria in a mouse model of malaria infection, although no significant difference in parasitaemia was recorded and the mechanism linking the two is still unclear (Robert, *et al* 2008). Other studies, also in mice, have noted a correlation between malaria infection and outcome in hypoglycaemia and hyperinsulinemia models (Elased and Playfair 1994), as well as with mice under calorie restriction (Mancio-Silva, *et al* 2017). The latter study was found to be due to nutrient sensing by the parasites and subsequent adjustment of multiplication rates through changes in gene expression levels according to nutrient availability. Finally, in humans, assessment of clinical malaria cases in Nigeria noted a correlation between malaria susceptibility and serum cholesterol levels (Chukwuocha and Eke 2011) though the power of the study was relatively low.

Given the implied linkages between diet, cholesterol levels and malaria and the clear role cholesterol plays in defining membrane properties of cells, we set out to test how elevated cholesterol levels affect erythrocyte biophysical properties and susceptibility to malaria parasite infection using both an *in vitro* human *P. falciparum* and murine *P. berghei* model.

## RESULTS

### Artificial incorporation of membrane cholesterol inhibits *P. falciparum* merozoite invasion

Based on our previous findings demonstrating that a reduction in the erythrocyte bending modulus resulted in an increased merozoite invasion efficiency (Koch, *et al* 2017), we hypothesised that stiffening the erythrocyte membrane by increasing cholesterol content would result in a lower merozoite invasion efficiency. Erythrocytes were incubated in media supplemented with a range of cholesterol concentrations, however, no change in the red cell bending modulus was found (Fig 1A-D). Methyl-β-cyclodextrin (MβCD) is a compound frequently used to incorporate additional cholesterol into cell membranes. Because of its high affinity for cholesterol, MβCD can be used directly to extract cholesterol from cell membranes or it can be coupled with cholesterol prior to addition to cells in order to increase cholesterol packing in the target cell membrane (Zidovetzki and Levitan 2007). Thus, in an attempt to increase erythrocyte cholesterol content, red cells were incubated with MβCD coupled cholesterol. As expected, a highly significant increase in erythrocyte bending modulus was observed, whilst no significant increase in membrane tension (a measure of cytoskeletal effects (Koch, *et al* 2017) was found at this concentration (Fig 1E-F). To confirm that additional cholesterol had been incorporated into the cell membranes, cells were washed, lipid membrane extracted, purified, dried and resolubilised to measure cholesterol content. A concentration dependent increase in membrane cholesterol was observed in erythrocytes preincubated with MβCD coupled cholesterol (Fig 1 G). To ensure that experimental samples fell within a linear range, a cholesterol fluorometric standard curve was set up (Fig 1H). To test whether a cholesterol-dependent increase in bending modulus is negatively correlated with merozoite invasion efficiency, erythrocytes were exposed to various concentrations of MβCD-coupled cholesterol, washed and then incubated with purified merozoites added for 30 minutes. Invasion efficiency was quantified using flow cytometry and showed a clear linear relationship between cholesterol-dependent increase in bending modulus and merozoite invasion (Fig 2A-C).

**Fig 1.**
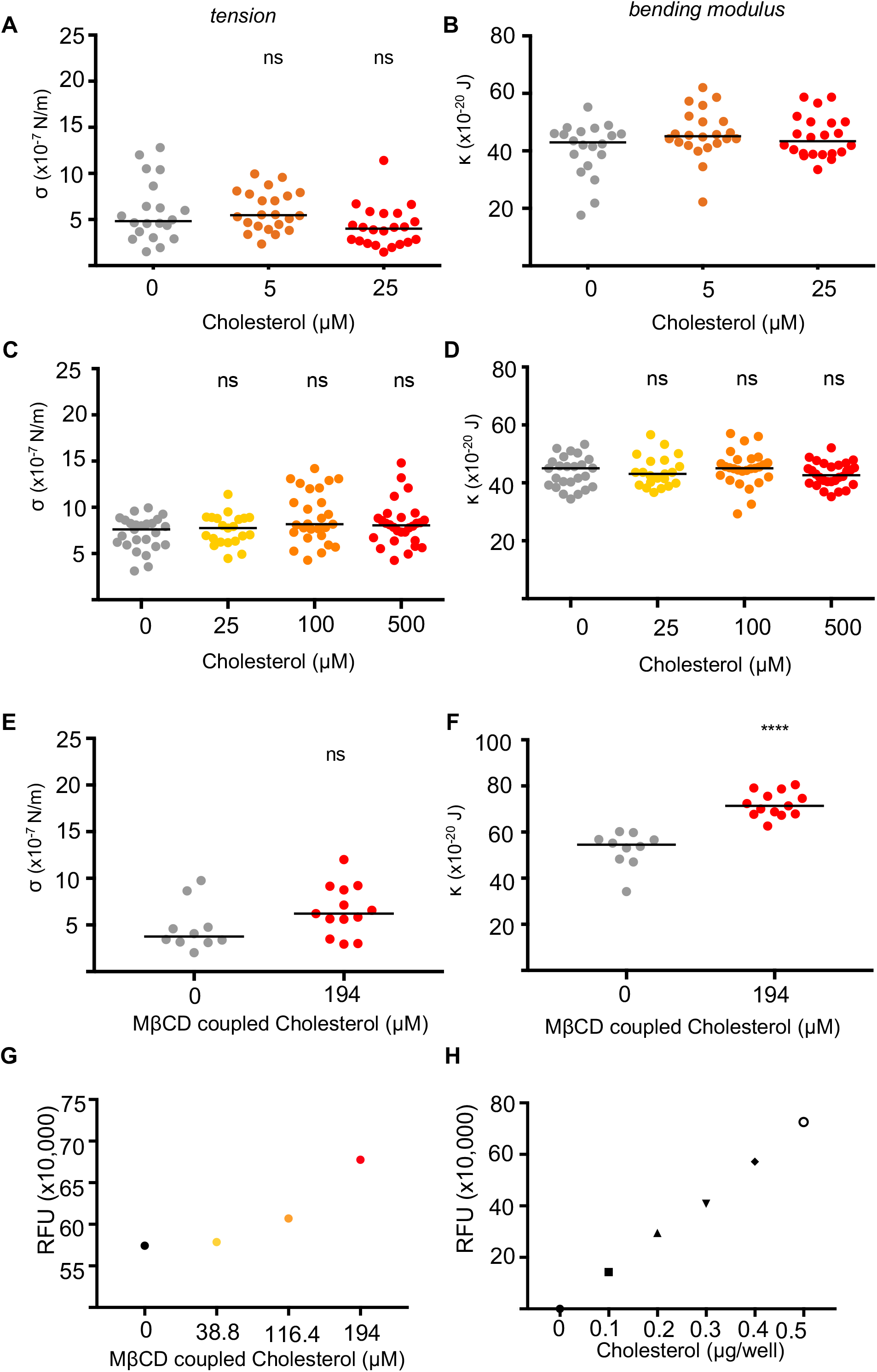
Incubating red cells with cyclodextrin coupled cholesterol leads to incorporation of cholesterol into red cell membranes and increased red cell bending modulus. Repeated attempts to increase red cell cholesterol by incubating cells in media supplemented with cholesterol only did not result in changes to red cell biophysical properties. Summary of (A & C) tension (σ) and (B & D) bending modulus (κ) values of red cells from different donors pre-treated with a wide range of cholesterol concentrations. Pre-treatment of red cells with cholesterol coupled to cyclodextrin similarly did not affect red cell (E) tension (σ) but significantly increased red cell (F) bending moduli (κ). Each circle represents data from a single cell, and the solid line represents the median. To quantify the red cell cholesterol, first a cholesterol fluorometric standard curve was set up to show the linear range of the cholesterol quantitation assay. Red cells were incubated with increasing amounts of cyclodextrin coupled cholesterol, washed and lipid membranes purified prior to cholesterol quantification. (H) A concentration dependent increase in the cholesterol content was found in erythrocytes that had been incubated with cyclodextrin coupled cholesterol. P values comparing the treatment versus control flicker spectroscopy data were calculated using the Mann–Whitney test (ns = not significant; **** p < 0.0001).

**Fig 2.**
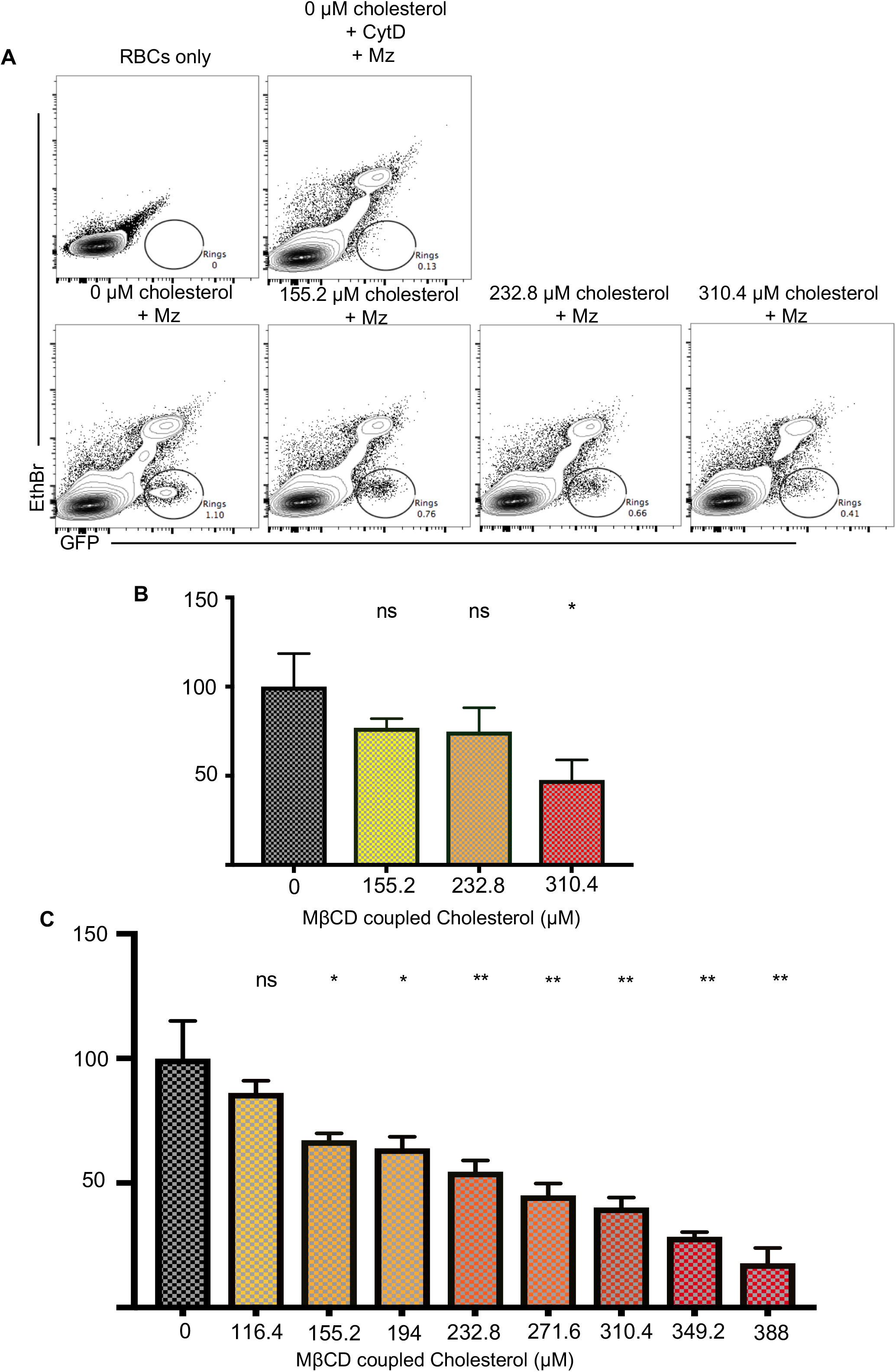
Increasing the red cell membrane cholesterol content reduces *P. falciparum* invasion efficiency. (A) Representative Flow Cytometry profiles highlighting infected erythrocyte populations. (B) Quantification of *P. falciparum* invasion efficiency into cyclodextrin-cholesterol pretreated red cells. Values represent mean and standard deviation of triplicate wells. This effect on erythrocyte invasion was reproducible across different blood donors (C). P values comparing the treatment versus control were calculated using the Student’s t test (* p < 0.05; ** p < 0.01).

### Establishing an *in vitro* merozoite invasion assay for *P. berghei*

Having established a clear link between cholesterol content, erythrocyte bending modulus and *P. falciparum* merozoite invasion efficiency, we next sought to explore whether increased serum cholesterol *in vivo* might partially protect the host from merozoite invasion using the murine malaria model *P. berghei.* Before we could commence study of cholesterol effects, we first had to develop a workflow that can isolate invasion events in the mouse malaria model. To date, study of merozoite invasion in *P. berghei* (as opposed to infection *in vivo)* has lagged behind that of *P. falciparum* (Boyle, *et al* 2010) and *P. knowlesi* (Lyth, *et al* 2018) largely because, as an *in vivo* model, there are few *in vitro* tools for its long-term propagation and therefore ability to isolate synchronous invasion events. Thus, to enable direct measurement of invasion (as opposed to growth) we sought to establish a robust workflow for merozoite isolation and invasion assessment of murine erythrocytes using *P. berghei in vitro.* Quantifying murine erythrocyte parasitemia usually uses flow-cytometry based on the nuclear staining of infected cells. The presence of different nucleated cell populations in mouse blood (Howell-Jolly bodies, red cell progenitors as well as leukocytes) necessitates use of a combination of antibodies and stains to allow identification of infected erythrocytes (Lelliott, *et al* 2014). A further complication arises because of the lack of synchronicity of the parasites *in vivo,* with invasion occurring over a course of several hours – which can vary between individual animals – making comparison difficult. To overcome these limitations, we adapted the *in vitro* method for quantifying *P. falciparum* merozoite invasion (Boyle, *et al* 2010) to *P. berghei* (Fig 3). Using GFP-positive *P. berghei* we could ensure the isolation of synchronous parasites that, on maturation, can be magnet separated and 1.2 μm filtered to release individual infectious merozoites. Using a 96 well-plate format a fixed volume of free merozoites were added to 10^6^ erythrocytes, allowing for 30 minutes invasion at 37 °C while shaking. Staining samples with Ethidium bromide, samples could then be analysed by flow cytometry. This workflow enabled an invaded parasite population, gated on being GFP-positive and low EtBr staining (as the dye does not stain the intra-erythrocytic parasite as efficiently (Wilson, *et al* 2013) to be determined, accurately isolating our ability to measure parasite invasion (Fig 3).

**Fig 3.**
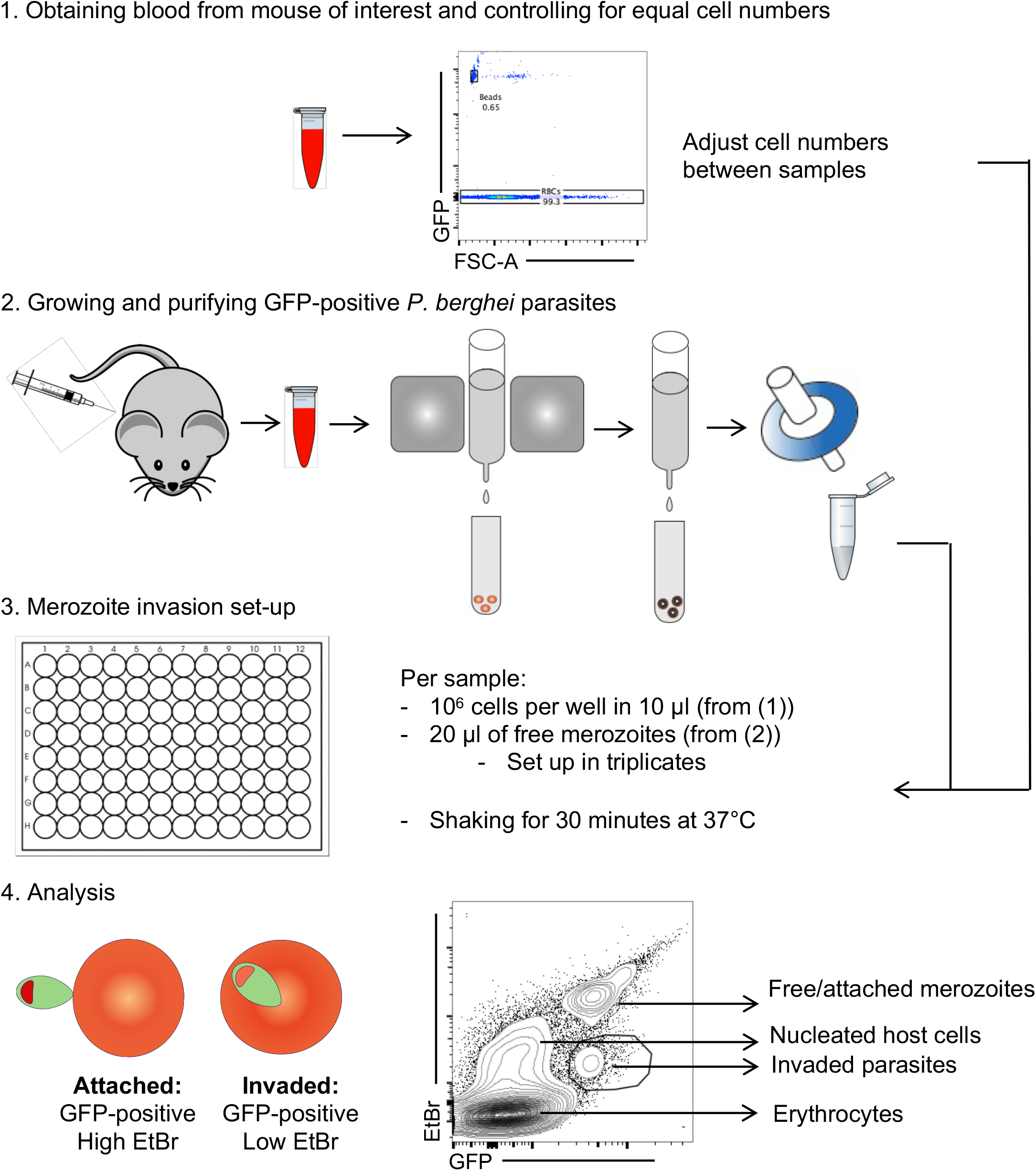
Workflow of the *P. berghei* merozoite assay. Firstly, the blood from the mouse (or mice) of interest is harvested the day prior to the invasion assay either by tail-bleed or cardiac puncture. For the assay to be quantitative, the number of cells need to be kept equal between samples obtained from different mice. This is done using a flow cytometric counting assay. Secondly, GFP-positive *P. berghei* parasites are injected and grown in mice. At the appropriate parasitemia, the blood is harvested by cardiac puncture one day prior to the invasion assay and incubated at 37C under low oxygen conditions until the parasites have developed into mature schizonts (12-24h). The mature parasites are then separated from uninfected blood using a MACS magnetic cell separator, and passed through a 1.2 micrometer filter to rupture the parasitophorous vacuolar membrane and release individual merozoites (Boyle, *et al* 2010). Next, in a 96 well plate, three wells per mouse, each with 1 x 10^6^ blood cells, are set up before a fixed volume of free merozoites is added to each well. Invasion is allowed to occur for 30 minutes at 37°C while shaking. Finally, the samples are stained with EtBr and analysed by flow cytometry. The invaded parasite population is gated on based on their GFP-positive, low EtBr staining, as the dye does not stain the intra-erythrocytic parasite as efficiently (Wilson, *et al* 2013).

### *P. berghei* merozoite invasion efficiency is not affected by increased serum cholesterol in a mouse model of hypercholesterolemia

To test whether hypercholesterolemia does indeed stiffen the erythrocyte membrane and provides some protection against *P. berghei* merozoite invasion, we sought an appropriate mouse model of hypercholesterolemia. Unlike humans, mice do not efficiently absorb excess cholesterol through diet and are therefore protected from hypercholesterolemia (Carter, *et al* 1997) without perturbation of cholesterol metabolism. Several genetically modified mouse models have been developed to overcome this issue; by far the most commonly used are LDL receptor knockout (LDL-R^−/−^) and ApoE knockout (ApoE^−/−^) strains (Zadelaar, *et al* 2007). Perturbations of either pathway disrupt reverse cholesterol transport leading to excess LDL cholesterol in the circulation.

### *P. berghei* merozoite invasion was tested in the hypercholesterolemia susceptible mouse strains

LDL-R^−/−^. Six mice were used, three placed on a standard diet with the other three placed on a high fat diet for 8 weeks before blood was harvested by cardiac puncture and used for quantitative merozoite invasion assays. Using two independent parasite cultures on separate days, no significant difference was found in merozoite invasion efficiency between the standard and high-fat diet groups (Fig 4 A-C, D-E). Thus, although the sample size was small, no correlation was found between merozoite invasion efficiency and total plasma cholesterol (Fig 4F).

**Fig 4.**
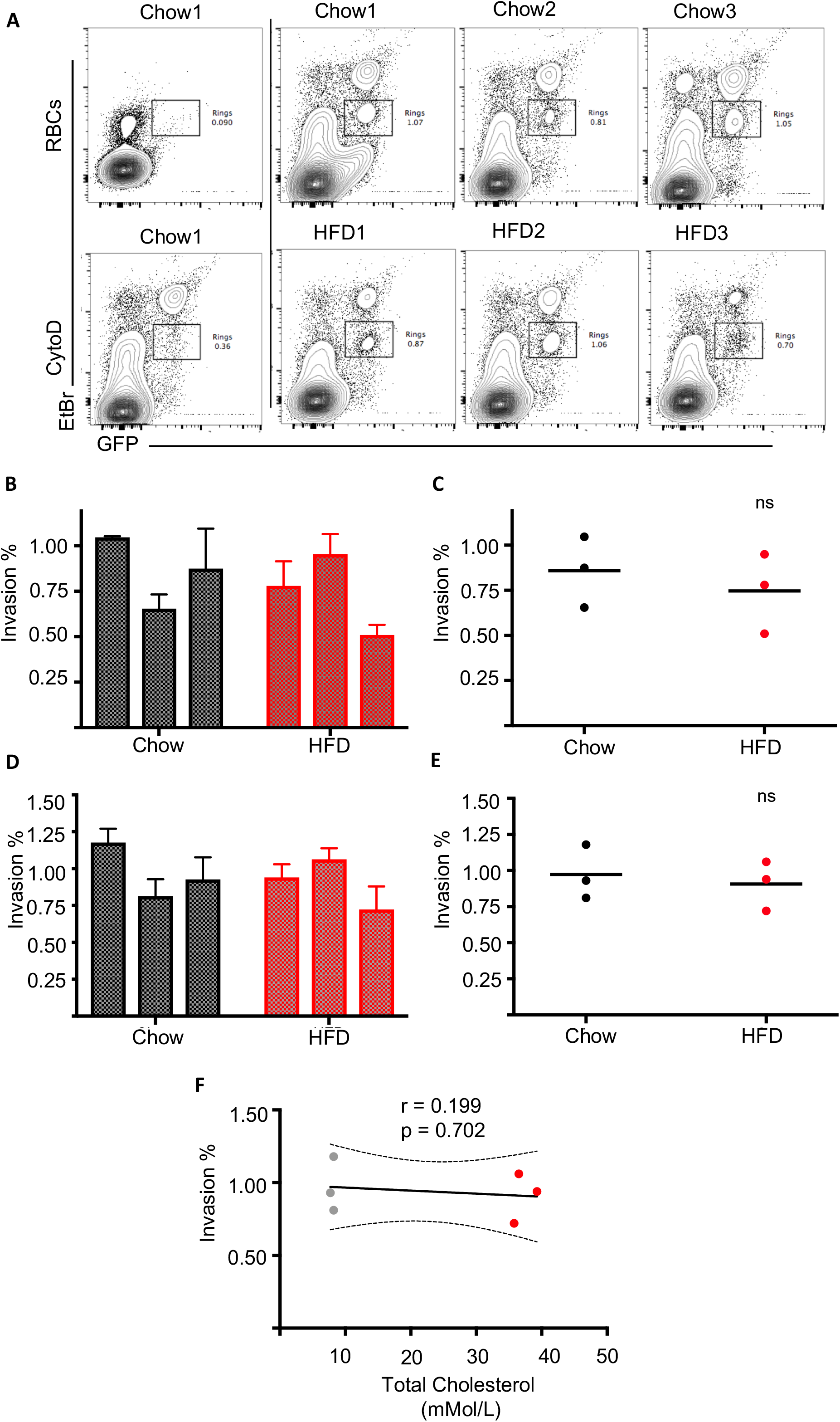
*P. berghei* invasion rates into red cells from LDL-R-/- mice do not differ significantly between chow fed and high fat diets. Invasion rates obtained from Chow and High-Fat Diet (HFD) fed LDL-R-/-. (A) Representative flow cytometry profiles highlighting infected red cell populations obtained from mice fed the standard Chow diet (Chow) and high fat diet (HFD). A blood sample in the absence of merozoites and a sample with merozoites and CytD are shown as gating controls. (B & D) Quantification of *P. berghei* invasion assays carried out on two separate days with newly harvested parasites obtained from different mice. Values represent mean and standard deviation of triplicate wells. The invasion percentages summarised per group are shown for both experiments in (C & E). Each circle represents the average invasion percentage per sample, line represents mean. No significant differences were found in erythrocyte invasion efficiency between Chow and HFD groups. The correlation between red cell invasion and total plasma cholesterol is shown in (F) along with the Pearson correlation coefficient (r), p value and a linear regression line modeling the correlation with 95 % confidence intervals (CI). P values comparing *P. berghei* invasion into red cells from HFD or Chow fed mice were calculated using the Student’s t test (ns = not significant).

### Erythrocytes from mice with high serum cholesterol do not show altered biophysical properties

To explore why serum cholesterol did not correlate with invasion efficiency, as it had *In vitro* with *P. falciparum* merozoites into erythrocytes with artificially elevated membrane cholesterol, we explored whether high fat diet resulted in increased serum cholesterol levels and stiffer erythrocyte membranes in LDL-R^−/−^ mice. Plasma cholesterol was quantified and the erythrocyte bending modulus was measured using flicker spectroscopy. The plasma cholesterol and triglyceride levels of LDL-R ^−/−^ mice are summarised in Table 1. Plasma cholesterol levels of mice fed the high fat diet were significantly higher (p<0.0001) than mice fed the standard diet as expected. Both HDL and LDL levels were elevated, however the biggest increase was observed in the LDL-cholesterol levels. A minimum of 20 erythrocytes from each Chow fed and HFD LDL-R^−/−^ mice were then analysed using flicker spectroscopy. Tension and bending modulus values of individual erythrocytes are summarised in Fig 5 A-B. Notably, the median tension levels of the LDL-R^−/−^ mice were found to be comparable across all samples, ranging between 4.6 and 5.4×10^-7^ N/m except for Mouse C2 for which the median tension is considerably higher at 7.4×10^-7^ N/m (Fig 5C). The reason for the higher erythrocyte tension in this mouse is not known. The median bending modulus values were also found to be comparable across the samples, ranging from 34.2 to 38.3 (Fig 5 D). Thus, no significant correlation was found between plasma cholesterol levels and erythrocyte bending modulus. This suggests that the large increase in the plasma cholesterol levels in the HFD fed mice does not significantly impact on the erythrocyte membrane cholesterol in mice.

**Fig 5.**
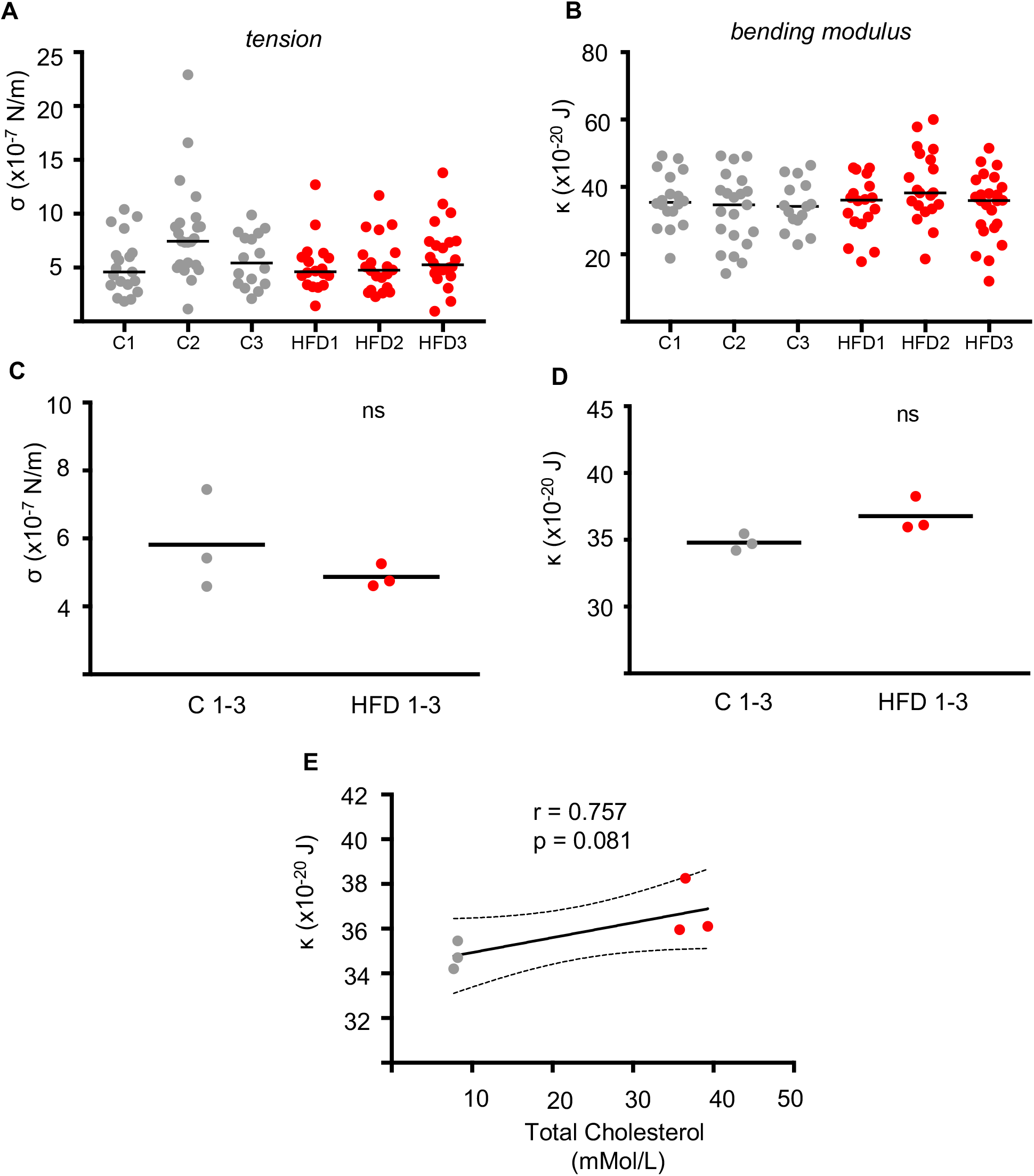
No significant changes in red cell biophysical properties were found between cells obtained from Chow or HFD-fed LDL-R(-/-) mice. Summary of tension (A) and bending modulus (B) values measured using flicker spectroscopy. Each circle represents data from a single cell, the solid line represents the median. Inter-group comparison of Chow and HFD erythrocyte tension (C) and bending modulus values (D) revealed no significant differences between the groups. Each dot represents the median tension and bending modulus obtained in (A-B), solid line represents the mean. The correlation between bending modulus (κ) and total plasma cholesterol is shown in (E) along with the Pearson correlation coefficient (r), p value and a linear regression line modeling the correlation with 95 % confidence intervals (CI). P values comparing the HFD versus control were calculated using the Student’s t test (ns = not significant).

**Table 1.** Plasma cholesterol quantification of LDL-R-/- mice

| Sample | Cholesterol<br>(mmol/L) | Triglycerides<br>(mmol/L) | HDL<br>(mmol/L) | LDL<br>(mmol/L) |
| --- | --- | --- | --- | --- |
| Chow 1 | 8.2 | 1.6 | 1.65 | 5.8 |
| Chow 2 | 8.2 | 1.7 | 1.23 | 6.2 |
| Chow 3 | 7.7 | 2.4 | 2.56 | 4.0 |
| HFD 1 | 39.3 | 4.5 | 2.33 | 34.9 |
| HFD 2 | 36.5 | 4.5 | 2.44 | 32.0 |
| HFD 3 | 35.8 | 4.5 | 2.43 | 31.3 |

### *P. falciparum* merozoite invasion efficiency is not affected by increased serum cholesterol in patients with hypercholesterolemia

Since metabolism in mice differs from humans in a number of ways as discussed above, human blood samples from patients with a range of cholesterol levels were collected and tested towards linking plasma serum cholesterol and merozoite invasion efficiency. Blood samples from patients with normal cholesterol levels (up to 5 mMol/L) were compared with samples from patients with heterozygous familial hypercholesterolemia (elevated levels: between 5 and 10 mMol/L) as well as samples from patients with homozygous familial hypercholesterolemia (severely elevated cholesterol levels: above 10mMol/L). Strikingly, and mimicking that found with mouse models, the level of serum cholesterol again had no significant impact on *P. falciparum* merozoite invasion efficiency (Fig 6 A-C).

**Fig 6.**
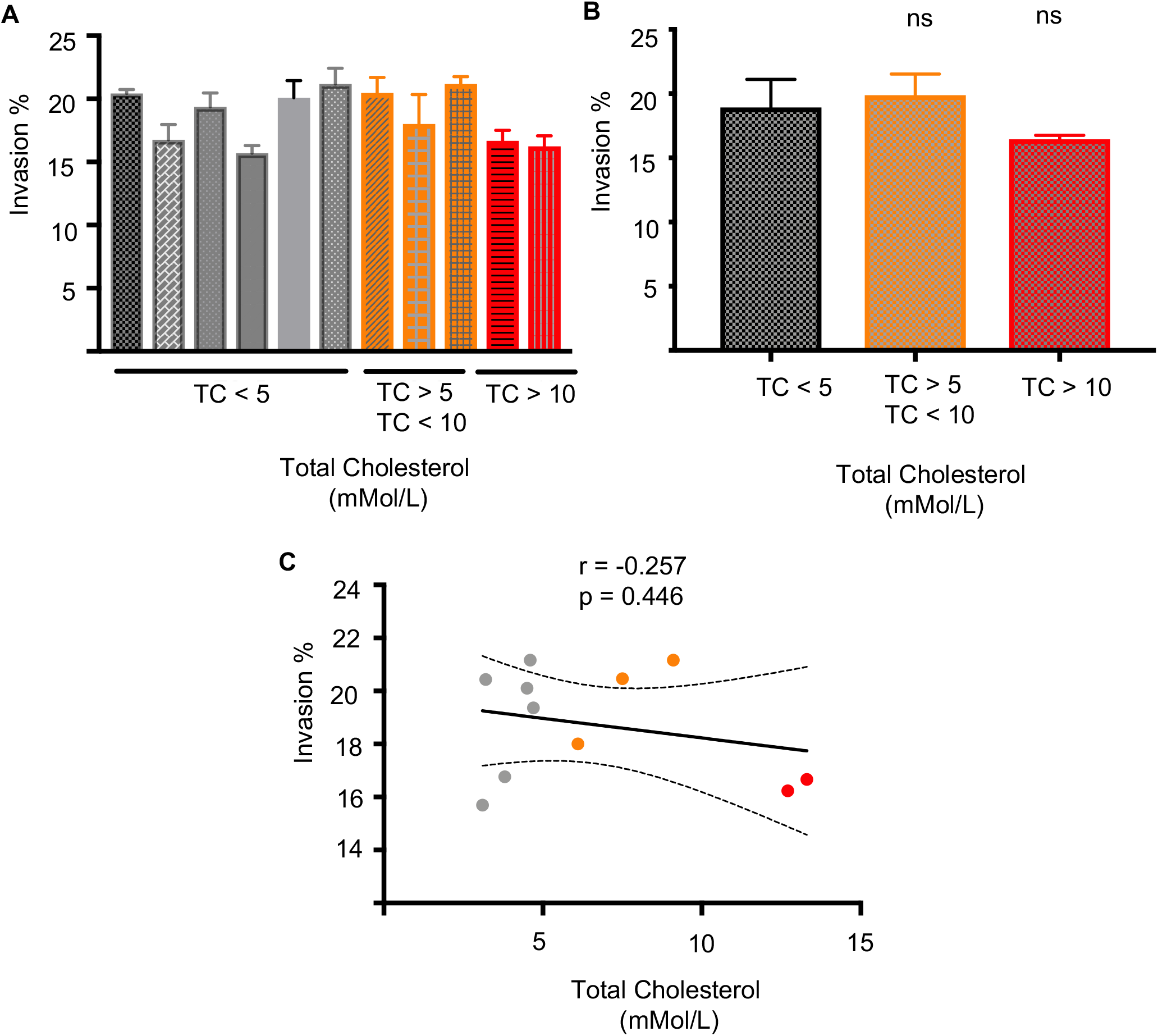
*P. falciparum* invasion rates into red cells obtained from patients with varying total plasma cholesterol (TC) levels do not differ significantly. (A) Quantification of *P. falciparum* invasion rates into red cells from patients with low TC (<5), elevated TC (>5, <10) and severely elevated TC (>10). Values represent mean and standard deviation of triplicate wells. The invasion percentages summarised per group are shown for both experiments in (B). The correlation between red cell invasion and total plasma cholesterol is shown in (C) along with the pearson correlation coefficient (r), p value and a linear regression line modeling the correlation with 95 % confidence intervals (CI). P values comparing *P. falciparum* invasion into red cells from low, elevated and highly elevated TC samples were calculated using the Student’s t test (ns = not significant).

### Erythrocytes from patients with high serum cholesterol do not show altered biophysical properties or cholesterol content

To parallel our studies with murine erythrocytes, we investigated the effects of hypercholesterolemia on erythrocyte biophysical properties, tension and bending modulus values for individual red cells from each patient sample (Fig 7A-B), with group summaries shown in Fig 7 C-D. Again, paralleling mouse studies, no significant correlation was found between total serum cholesterol and red cell bending moduli (Fig 7E). Finally, to investigate the relationship between plasma cholesterol and erythrocyte membrane cholesterol, red cell membranes from each clinical sample were dried, purified and quantified (Fig 8 A-C). No significant correlation was found between erythrocyte membrane cholesterol and plasma serum cholesterol (Fig 8D). Thus, despite serum cholesterol levels above 10mMol/L, it appears that human erythrocytes (and likely murine erythrocytes) are naturally buffered from incorporation of excess cholesterol. This would suggest that whilst cholesterol content of the erythrocyte is a direct correlate of malaria parasite invasion efficiency, this is not affected by plasma cholesterol levels.

**Fig 7.**
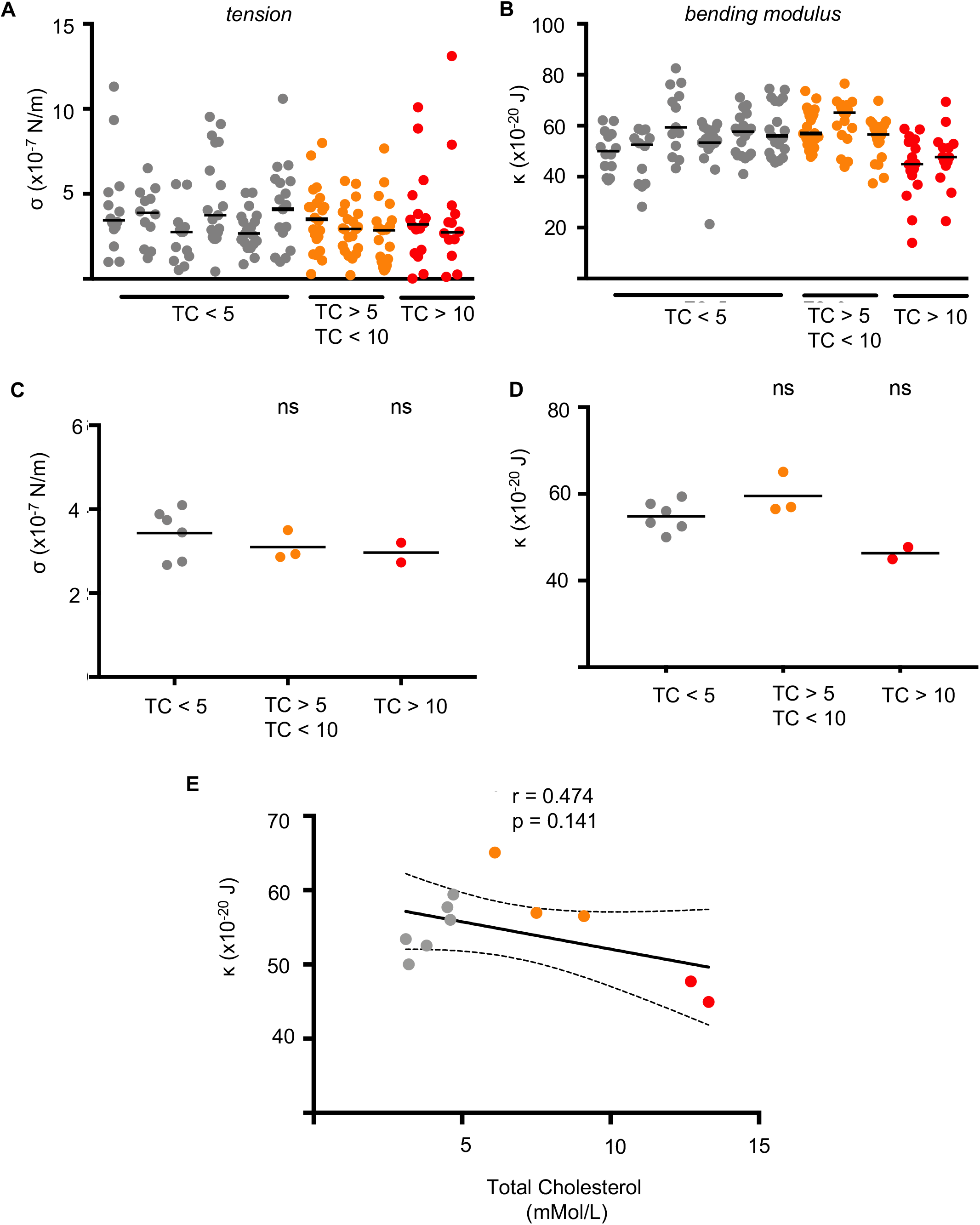
No significant changes in red cell biophysical properties were found between cells obtained from patients with varying total plasma cholesterol (TC) levels. Summary of tension (A) and bending modulus (B) values measured using flicker spectroscopy. Each circle represents data from a single cell, the solid line represents the median. Inter-group comparison of low TC, elevated and highly elevated TC red cell tension (C) and bending modulus values (D) revealed no significant differences between the groups. Each dot represents the median tension and bending modulus obtained in (A-B), solid line represents the mean. The correlation between bending modulus (κ) and total plasma cholesterol is shown in (E) along with the Pearson correlation coefficient (r), p value and a linear regression line modeling the correlation with 95 % confidence intervals (CI). P values comparing the HFD versus control were calculated using the Student’s t test.

**Fig 8.**
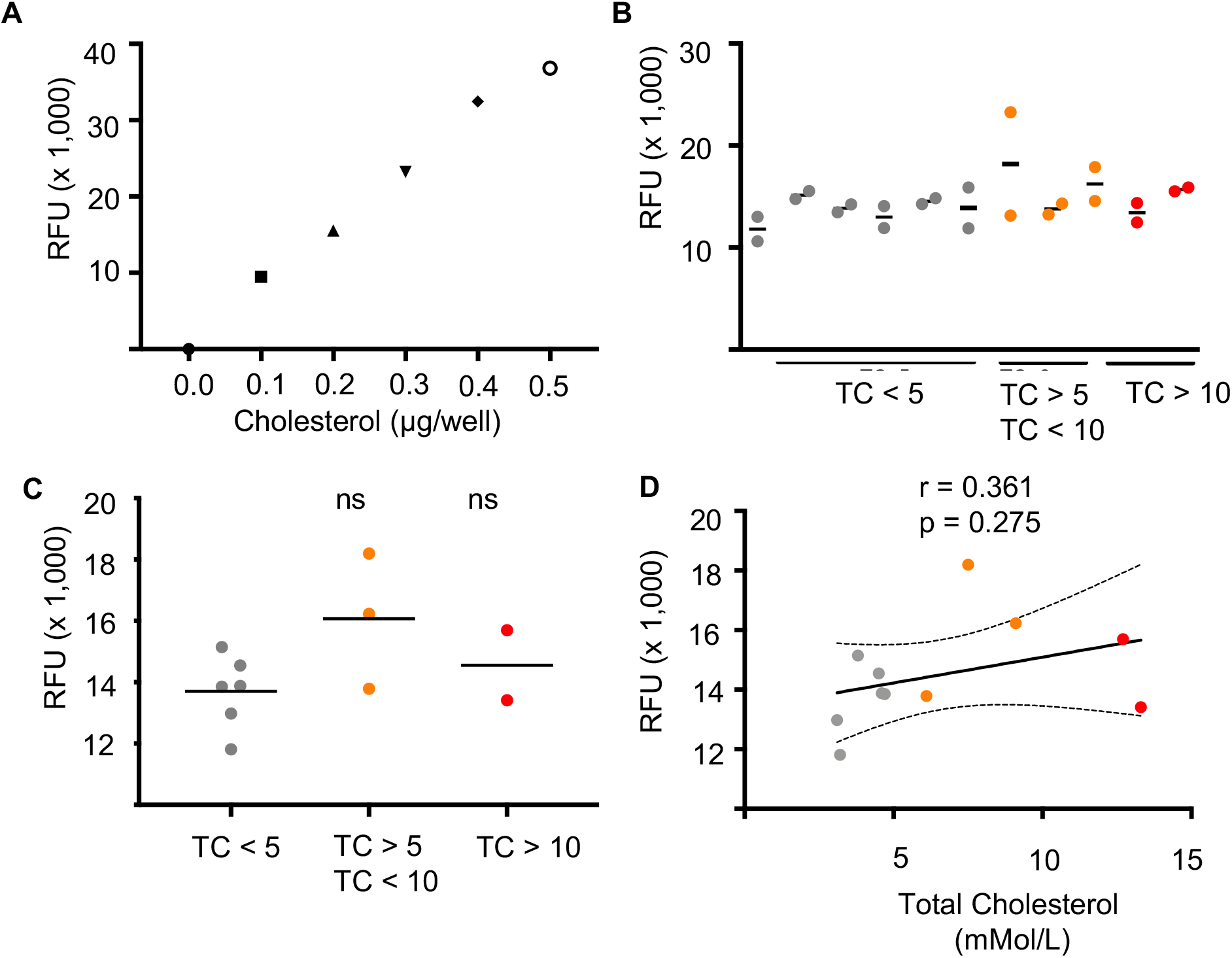
Plasma cholesterol is not correlated with red cell membrane cholesterol. (A) Fluorometric cholesterol standard curve (B) Red cell membranes from patients with low, elevated and highly elevated total plasma cholesterol were extracted and dried before being quantified using the Cholesterol Quantitation kit (Sigma). (C) Inter-group comparison of Chow and HFD cholesterol measurements revealed no significant differences between the groups, solid line represents the mean. The correlation between bending modulus (κ) and total plasma cholesterol is shown in (E) along with the pearson correlation coefficient (r), p value and a linear regression line modeling the correlation with 95 % confidence intervals (CI). P values comparing membrane cholesterol between patient samples were calculated using the Student’s t test. RFU = Relative Fluorescence Units.

## DISCUSSION

By artificially manipulating erythrocyte cholesterol content, we confirmed the importance of red cell biophysics in merozoite invasion showing that the erythrocyte bending modulus (a measure of membrane deformability) is markedly affected by membrane cholesterol content, and that this is negatively correlated with merozoite invasion efficiency. To investigate whether medical conditions associated with increased cholesterol levels lead to stiffer red cells and an associated reduction in merozoite invasion efficiency, we quantified the biophysical properties of erythrocytes obtained from a murine hypercholesterolemia model as well as from human clinical samples with reported hypercholesterolaemia and subsequently used these in quantitative merozoite invasion assays. In both the murine model of hypercholesterolemia and clinical samples of patients with elevated serum cholesterol, these levels did not correlate with changes in erythrocyte cholesterol content and hypercholesterolemic blood samples showed comparable malaria parasite infection rates to matched controls.

Due to the difficulty in obtaining large numbers of fresh blood samples from hypercholesterolemic patients not on statins, it is possible that very subtle effects on red cell cholesterol content may not be found with the given sample size. Therefore, we decided to investigate any potential effects on erythrocyte membrane cholesterol content within a clinical setting as well as using a mouse model, and obtained consistent results across the two. We also included blood samples of patients with homozygous familial hypercholesterolemia, a rare condition which leads to extremely elevated levels of serum cholesterol. Even in these rare cases, red cell membrane cholesterol did not differ significantly from samples obtained from healthy patients. This implies cholesterol content in the erythrocyte is tightly regulated and not significantly affected by serum cholesterol content.

The lack of correlation between plasma and red cell membrane cholesterol was surprising considering that a similar study carried out in two animal models of hypercholesterolemia found both plasma and membrane cholesterol levels responding to diet and medication (Sengupta and Ghosh 2011, Tziakas, *et al* 2007), suggesting that erythrocyte membrane cholesterol is dynamic and modifiable under certain conditions. In humans, the picture is not as clear, while both diet (Cazzola, *et al* 2011) and lipid lowering drugs (Martinez, *et al* 1996) have been shown to affect red cell cholesterol content, other studies report no direct correlation between plasma and red cell membrane cholesterol content (Tziakas, *et al* 2007). Importantly, how erythrocyte cholesterol content is regulated is not well understood, since erythrocytes neither produce cholesterol *de novo* nor contain receptors, such as LDL-R, which allow uptake of lipoprotein molecules (D’Alessandro, *et al* 2010). The most likely mechanism for the induced changes in red cell cholesterol content is a dynamic exchange between plasma and membrane cholesterol (Gold and Phillips 1990, Quarfordt and Hilderman 1970), with the rate of exchange likely being influenced by the structure and composition of the cell (Gold and Phillips 1990).

The work presented here suggests that a diet induced increase in plasma cholesterol does not significantly affect erythrocyte membrane cholesterol and these cells are therefore not protected from merozoite invasion. Though diet and serum cholesterol do not appear to be the best predictors of erythrocyte cholesterol content, we believe other metabolic causes of erythrocyte biophysical variability caused by lipid content is possible. Future work focusing on sampling and identifying abnormal red cell biophysical properties in malaria endemic regions, we predict, could still find new protective polymorphisms against malaria disease.

## ACKNOWLEDGEMENTS

We thank Jane Srivastava for assistance with flow cytometry (Imperial College Flow Cytometry Facility), Kate Wright and Fernando Sanchez-Roman Teran for assistance with *P. falciparum* merozoites and Jason L Johnson for generously providing additional mouse models (eventually not used in this study). Tissue sample collection was supported by the National Institute for Health Research (NIHR) Biomedical Research Centre based at Imperial College Healthcare NHS Trust and Imperial College London. M.K. is supported by a PhD scholarship from the UK Medical Research Council (MR/K501281/1). B. J. is supported by an NIHR Clinical Lectureship. Research was directly supported by an Investigator Award from Wellcome (100993/Z/13/Z J.B.). The views expressed are those of the authors and not necessarily those of the NHS, the NIHR or the Department of Health.

## METHODS

### Human blood samples and serum cholesterol measurements

Human erythrocytes (O+, male) for parasite invasion work were obtained from the NHS Blood Transfusion Service. Approval for collection of clinical human blood samples was granted via the Imperial College Healthcare Tissue Bank, National Research Ethics approval number 17/WA/0161, project ID R18015. Blood samples were collected from hypercholesterolaemic patients attending the Imperial College London NHS lipid clinic, and from severely hypercholesterolaemic patients undergoing lipoprotein apheresis as a treatment for homozygous hypercholesterolaemia. EDTA whole blood was collected for parasite assays and parallel measurement of serum cholesterol was performed (Abbott Architect assay, North West London Pathology Blood Sciences Laboratories). Anonymised, normocholesterolaemic EDTA whole blood samples, originating from a primary care setting, were obtained from the hospital blood sciences laboratory. No samples from patients with haemoglobinopathy were included.

### Hypercholesterolaemia mouse model and serum cholesterol measurement

C57/B6 *Ldlr^−/−^* mice were obtain directly from the Jackson Laboratory (https://www.jax.org/). The mice were fed either on normal chow (SAFE diet 105) or Western High Fat (Dietex, FAT 21%, Cholesterol 0.15%) diets for 8 weeks. Total cholesterol and HDL cholesterol were measured using an enzymatic method in a Siemens Dimensions RxL analyser, following manufacturer’s instructions.

### Cyclodextrin-complexed cholesterol

Cholesterol (Sigma-Aldrich) was made up in Ethanol at a concentration of 15 mg/ml. A 5% methyl-bcyclodextrin (MbCD) stock was made up in MQ-water and heated up on a hotplate stirrer set to 80 _C. 4 x 10 μl aliquots of the 15 mg/ml cholesterol stock was added to 400 μl of the MbCD stock on the stirrer, leaving 10 minutes between each aliquot. The MbCD-cholesterol mixture was left stirring on the hotplate for one hour (to evaporate the ethanol and allow complex formation) and either used immediately or stored at -20 C. The final concentration achieved is 38.8 mM MbCD and 3.8 mM cholesterol in MQ-water. Erythrocytes were incubated with MbCD-cholesterol complexes for 30 minutes at room temperature while shaking, with concentrations of up to 3.88 mM MbCD – 388 μM cholesterol. Following incubation, erythrocytes were spun down at 800 x g and resuspended into fresh RPMI media before use for further experiments.

### Erythrocyte membrane cholesterol measurements

Membrane cholesterol was quantified using a fluorometric Cholesterol Quantitation Kit (Sigma-Aldrich, UK) according to manufacturer instructions. In short, membranes were extracted with a chloroform-isopropanol-IGEPAL CA-630 solution and spun at 13,000 x g for 10 minutes, before the organic phase was transferred to a new tube and dried under nitrogen. Samples were put under vacuum for 30 minutes to remove any residual organic solvents before the lipid films were dissolved in Cholesterol Assay Buffer and vortexed until the mixture was homogenous. Fluorescence intensity was measured using a Tecan MPro 200 fluorescent plate reader (excitation: 535, emission: 587 nm).

### Parasite *in vitro* culture

*Plasmodium falciparum* parasites (D10-PHG (Wilson, *et al* 2010) were maintained in standard culture conditions (Trager and Jensen 1976). Parasites were grown in human O+ erythrocytes at 4% hematocrit in complete RPMI1640-HEPES media (Sigma-Aldrich) supplemented with 0.3% L-glutamine, 0.05% hypoxanthine, 0.025% gentamicin and 0.5% Albumax II (Life Technologies). Parasites were grown at 37 C in 1% O2, 5% CO_2_ in N_2_ and D10-PHG cultures were supplemented with 25 ng/mL pyrimethamine and 5 μg/mL blasticidin (Sigma-Aldrich)

### *P. falciparum* Merozoite Invasion Assays

The *P. falciparum* merozoite assay was carried out according to published protocols (Boyle, *et al* 2010, Zuccala, *et al* 2016). For a standard merozoite invasion assay, approximately 90 ml of 5% synchronised late stage D10-PHG (constitutively expressing green fluorescent protein (GFP)) parasites were magnetically isolated as described in (Boyle, *et al* 2010), incubated with 10 μM of the cysteine protease inhibitor l-transepoxysuccinyl-leucylamido-(4-guanidino)butane (E64, Sigma-Aldrich) for up to 6 hours under standard culture conditions until at least 50 % of schizonts had developed into parasitophorous vacuole enclosed membrane structures (PEMS) (Salmon, *et al* 2001). To obtain free merozoites, PEMS were centrifuged at 800 x g for five minutes (no brake) and resuspended into a minimum of 750 μl of RPMI. The concentrated PEMS-solution is then passed through a 1.2 μm Ministart syringe filter (Sartorius Stedim Biotech) to release individual merozoites.

For quantifying erythrocyte invasion rates, 25 μl of the filtered merozoite solution was added to a prepared erythrocyte suspension (10 μl of 1.5 % haematocrit) in a 96 well plate and incubated at 37 C on a shaker (300 – 500 RPM) for 20 minutes. An additional sample containing an invasion inhibitor (typically 100 nM cytochalasin D) was included in each experiment and used for accurate gating of the invaded population. Parasites were stained with 100 μl of 5 μg/ml Ethidium Bromide (EtBr) for 10 minutes at room temperature, washed twice in 100 μl phosphate buffered saline (PBS), and resuspended into 60 μl. Invasion was quantified by flow cytometry, acquiring a total of 100,000 events per well using a BD Fortessa flow cytometer equipped with a high-throughput plate reader. Cells were excited at 488 nm and emission read between 515 and 545 nm for GFP positivity and between 600 and 620 for Ethidium Bromide positivity. New ring-stage parasites were quantified based on their GFPpositive, EtBr-low staining profile (Zuccala, *et al* 2016). Data analysis was carried out in FlowJo v10 by gating on cells based on the side-scatter to forward-scatter profile (i.e. gating out debris), selecting single cells based on the forward-scatter width to area ratio and finally GFP and EtBr profile. Data was visualised in Prism v7 (GraphPad).

### *P. berghei* Merozoite Assays

Female Theiler’s Original (TO) mice, 8-10 weeks of age (Envigo, UK) were injected with 200 μl glycerol stocks of blood infected with 3 % *P. berghei* ANKA 507 expressing GFP. When parasitemia reached approximately 3%, parasites were harvested by cardiac puncture and incubated overnight at 37 C in 30 ml complete RPMI-1640 media under low oxygen conditions.

The following day schizonts were purified using a magnetic separation column (MACS, Miltenyi Biotec), Purified schizonts were ruptured with a 1.2 μm Ministart syringe filter (Sartorius Stedim Biotech) to release individual merozoites and 25 μl merozoite solution was added to blood obtained from mice of interest (10 ul at 100,000 cells per μl, counted as described below) in a 96 well plate. The culture was incubated for 20 minutes on a shaker at 37°C. Cells were stained and quantified as per the *P. falciparum* merozoite assay ‘quantification of invasion by flow cytometry’ described above.

### Flow cytometry and bead counting

The counting bead (CountBright Absolute Cell Counting Beads, Thermo-Fisher) solution was made up of 400 μl PBS, 50 μl counting bead solution and 50 μl of 1 % haematocrit blood. To ensure equal number of beads across the samples, a mastermix of PBS-Counting beads was made up first, mixed well and separated into the number of samples required before the cell solution was added to the bead-mix. Analysis was carried out on a BD Fortessa flow cytometer. Cells were excited at 488 nm and emission read between 515 and 545 nm. 2,000 beads were acquired per sample. Using ratiometric analyses based on the number of beads and cells acquired, the number of cells per μl of sample can be calculated. Data analysis was carried out in FlowJo v10 by gating on cells based on based on the forward-scatter and GFP negativity and beads based on their forward scatter profile and GFP positivity.

### Flicker Spectroscopy

Flicker spectroscopy was carried out as previously described (Koch, *et al* 2017). Membrane oscillation recordings were taken on a Nikon Ti Microscope (objective lens: Nikon Plan Apo 100x 1.4 N.A oil immersion) using an OrcaFlash4.0 CMOS camera. Approximately 4,500 frames were recorded at a frame rate of 150 (+/− 10 frames per second (fps)) and an exposure time of 1 ms. Data analysis was carried out using a custom-built LabVIEW (National Instruments) program that detects and extracts membrane contours. Fourier transforming gives a fluctuation power spectrum of mean square mode amplitudes hh2(qx, y = 0) as a function of mode wavenumber (qx). From these data, the bending modulus (k) and tension (s) can be fitted using the following equation:

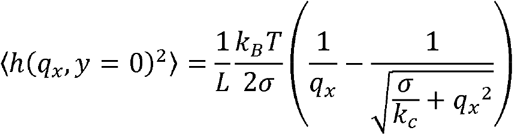

where k_B_ is the Boltzmann constant, T is temperature, and L is mean circumference of the cell contour (Koch, *et al* 2017).

## Supplementary Figures

**Supplementary Fig 1**. Red cell membrane cholesterol quantitation. (A) Fluorometric cholesterol standard curve (B) Red cell membranes from chow and HFD fed mice were extracted and dried before being quantified using the Cholesterol Quantitation kit (Sigma). (C) Inter-group comparison of Chow and HFD cholesterol measurements revealed no significant differences between the groups, solid line represents the mean. P values comparing HFD versus control were calculated using the Student’s t test. RFU = Relative Fluorescence Units.

